# Hybridization and rampant mitochondrial introgression among fire salamanders in peninsular Italy

**DOI:** 10.1101/258541

**Authors:** Roberta Bisconti, Daniele Porretta, Paola Arduino, Giuseppe Nascetti, Daniele Canestrelli

**Affiliations:** Tuscia University, Department of Ecological and Biological Science, Viterbo, 01100, Italy; University of Rome, “Sapienza”, Department of Environmental Biology, Rome, 00185, Italy

## Abstract

Discordance between mitochondrial and nuclear patterns of population genetic structure is providing key insights into the eco-evolutionary dynamics between and within species, and their assessment is highly relevant to biodiversity monitoring practices based on DNA barcoding approaches. Here, we investigate the population genetic structure of the fire salamander *Salamandra salamandra* in peninsular Italy. Both mitochondrial and nuclear markers clearly identified two main population groups. However, nuclear and mitochondrial zones of geographic transition between groups were located 600 km from one another. The overall pattern of genetic variation, together with morphological and fossil data, suggest that a rampant mitochondrial introgression triggered the observed mitonuclear discordance, following a post-glacial secondary contact between lineages. Moreover, at a shallower level of population structure, we observed evidence of asymmetric introgression of nuclear genes between two sub-groups in southern Italy. Our results clearly show the major role played by reticulate evolution in shaping the structure of *Salamandra salamandra* populations and, together with similar findings in other regions of the species’ range, contribute to identify the fire salamander as a particularly intriguing case to investigate the complexity of mechanisms triggering patterns of mitonuclear discordance in animals.

## Introduction

In principle, concordance between distinct phenotypic traits and/or genetic markers is a plausible expectation when analysing geographic patterns of biological diversity, above and below the species level. Derived from the shared history of individual characters of organisms, this expectation forms the basis for the extensive use of single markers (notably mitochondrial DNA) to draw inferences about the evolutionary history of populations and species^1^, as well as to assess their identity and geographic distribution using barcoding approaches^2, 3^. In practice, however, discordant patterns of variation among characters and among markers are not uncommon (e.g.^4–6^ among others). Several processes have been involved with the formation of such discordant patterns of variation, including various forms of selection, adaptive introgression, demographic disparities, hybrid zone movements, and sex-biased processes^7–10^. Consequently, discordance *per se* is emerging as a tremendous source of insights into the evolutionary process.

The drawbacks of using mitochondrial DNA (mtDNA) as the sole marker of genetic variation among and within populations are now widely acknowledged^11–13^, to the extent that a comparative approach using both mitochondrial (mtDNA) and nuclear (nuDNA) genetic data is becoming customary in phylogeographic investigations^14^. At the same time, however, after more than three decades when most phylogeographic studies adopted a single-marker approach (since^15^), there is now a huge amount of mtDNA data which needs to be complemented, in order to reach reliable conclusions about the evolutionary history and current genetic structure of populations. Furthermore, integrating mtDNA data with those drawn from the nuclear genome also has major practical implications. Indeed, mtDNA has been widely employed to revise classical taxonomy^16^, and to define conservation strategies for species and populations threatened by human impacts on the natural environment^17^. While in most cases such integration will hopefully lead to only slightly revisions of previous conclusions based on mtDNA data, there are compelling examples of the far reaching implications of a more integrative experimental approach to the study of population structure and history^7^.

The fire salamander *Salamandra salamandra* (Linnaeus, 1758) is a temperate amphibian largely distributed in Europe^18^. Owing to its remarkable phenotypic variation in traits encompassing external morphology, colour patterns, and reproductive strategy, the geographic variation and population structure of *S. salamandra* have been investigated in several portions of its range, using various combinations of phenotypic and genetic traits^19, 20^. As a consequence, the taxonomy of fire salamanders has been long discussed and repeatedly revised. Once regarded as a single highly polytypic species, the fire salamander group is now recognized as four distinct species, with more than 10 sub-species within *S. salamandra* alone^19^. At the same time, conflicting patterns of variation between traits have been frequently observed, and their analysis has formed the basis for intriguing insights into processes of reticulate evolution, range- change dynamics, and life-history traits evolution^21–23^. Consequently, the fire salamander is emerging as a compelling system for studying the contribution of multiple processes to the formation of intraspecific patterns of biological diversity^20^.

In this paper we investigate the population genetic structure of the fire salamander *S. salamandra* in peninsular Italy, a geographic area where instances of discordance between morphological traits and preliminary mtDNA data were previously observed^24^, without being reconciled into a population history, and whose underlying causes remain unclear. Based mostly on differences in colour patterns and body shape, fire salamander populations occurring along the Italian peninsula have long been attributed to the endemic subspecies *S. s. giglioli,* whereas populations from the pre-alpine and alpine areas have been attributed to the nominal subspecies *S. s.* salamandra^19, 25, 26^, with some intergradation between the two subspecies through the Liguria region (see Figure 1). On the other hand, a phylogeographic study of the mtDNA variation throughout the species range^24^ found mtDNA haplotypes typical of the northern subspecies *S. s. salamandra* occurring in south-central Italy, that is, several hundred kilometres to the south of the area of intergradation between the two subspecies, as outlined by phenotypic trait variation.

**Figure 1.**
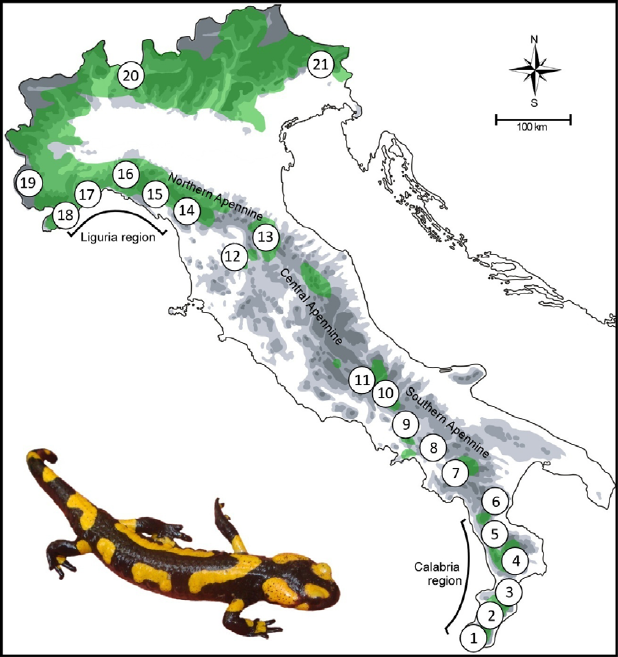
Geographic location of the 21 sampled populations of *S. salamandra,* and its approximate species’ distribution in Italy (light green; following^67^). Maps and diagrams were drawn using the software Canvas 11 (ACD Systems of America, Inc.). Photo: D. Canestrelli.

Here, using markers of both nuclear and mitochondrial genetic variation, in combination with phylogeographic and Bayesian population structure analytical tools, we aim to 1) assess the population genetic structure of the fire salamander throughout the Italian peninsula, 2) get a comprehensive understanding of the pattern of discordance observed between phenotypic traits and mtDNA data, and 3) shed light on its possible causes in these salamanders.

## Results

Nuclear genetic variation was investigated at level of 23 allozyme loci. Five loci (Mdhp-2, Mpi, Sod-1, Aat-2, and Ldh) were found monomorphic for the same allele in all the samples studied, while further three loci (Mdh-1, Mdh-2, Icdh-2) were found polymorphic at level of a single private allele, observed at low frequency (≤0.05). Allele frequencies observed at the remaining 15 loci are shown in Table 2. No statistically significant deviations were detected (at the 5% nominal level) from the expected Hardy-Weinberg (HW) equilibrium at each locus within populations, and from the expected genotypic linkage equilibria between pairs of loci within populations.

**Table 1.** Geographic location and sample size (n) of the 21 populations of *Salamandra salamandra* investigated in this study.

| Locality | Altitude<br>(m) | Latitude<br>(N) | Longitude<br>(E) | n<br>(allozymes) | n<br>(mtDNA) | mtDNA<br>haplotypes |
| --- | --- | --- | --- | --- | --- | --- |
| 1 Gambarie | 1310 | 38°10' | 15°50' | 16 | 17 | S1(4); S2(13) |
| 2 Carmelia | 1220 | 38°14' | 15°55' | 10 | 9 | S1(1); S2(6);<br>S3(1); S4(1) |
| 3 Serra S. Bruno | 805 | 38°35' | 16°20' | 7 | 6 | S2(6) |
| 4 Villaggio Mancuso | 520 | 39°01' | 16°35' | 11 | 11 | S5(8); S6(1);<br>S7(2); |
| 5 Fagnano Castello | 1090 | 39°33' | 16°01' | 3 | 3 | S8(2); S9(1) |
| 6 S. Severino Lucano | 890 | 40°04' | 16°06' | 7 | 11 | S10(7); S11(3);<br>S12(1) |
| 7 Laurino | 750 | 40°20' | 15°19' | 2 | 1 | S12(1) |
| 8 Giffoni | 770 | 40°46' | 14°55' | 8 | 8 | S13(2); N14(4);<br>N15(2) |
| 9 Cervinara | 550 | 41°01' | 14°36' | 11 | 10 | N14(10) |
| 10 Pescolanciano | 945 | 41°43' | 14°20' | 9 | 6 | N16(6) |
| 11 Val Fondillo | 930 | 41°42' | 14°21' | - | 8 | N16(8) |
| 12 Volpaia | 530 | 43°30' | 11°22' | 15 | - |  |
| 13 Camaldoli | 930 | 43°50' | 11°49' | 10 | 10 | N17(10) |
| 14 Cipollaio | 820 | 44°03' | 10°15' | 3 | 10 | N17(7); N18(3) |
| 15 Passo del Bracco | 610 | 44°15' | 9°34' | - | 5 | N17(5) |
| 16 Vallecaldà | 580 | 44°32' | 8°57' | 11 | 9 | N19(9) |
| 17 Manie | 350 | 44°12' | 8°22' | - | 3 | N20(3) |
| 18 Colle S. Bartolomeo | 320 | 43°59' | 7°57' | 10 | 8 | N17(8) |
| 19 Sampeyre | 950 | 44°34' | 7°11' | 8 | 8 | N17(7); N21(1) |
| 20 Banco | 660 | 46°00' | 8°50' | 8 | 5 | N22(5) |
| 21 Campone | 450 | 46°15' | 12°49' | 11 | 10 | N22(6); N23(2);<br>N24(1); N25(1) |

**Table 2.** Allele frequencies of the 15 allozyme loci found polymorphic among the 21 sampled populations of *Salamandra salamandra.*

| Population<br>Locus | 1 | 2 | 3 | 4 | 5 | 6 | 7 | 8 | 9 | 10 | 12 | 13 | 14 | 16 | 18 | 19 | 20 | 21 |
| --- | --- | --- | --- | --- | --- | --- | --- | --- | --- | --- | --- | --- | --- | --- | --- | --- | --- | --- |
| *G3pdh* |  |  |  |  |  |  |  |  |  |  |  |  |  |  |  |  |  |  |
| 85 | ---- | ---- | ---- | ---- | ---- | ---- | ---- | ---- | ---- | ---- | ---- | ---- | 0.17 | ---- | ---- | ---- | ---- | ---- |
| 100 | ---- | ---- | ---- | ---- | ---- | ---- | ---- | ---- | ---- | ---- | 0.25 | 0.20 | 0.50 | 1.00 | 1.00 | 1.00 | 1.00 | 1.00 |
| 115 | 1.00 | 1.00 | 1.00 | 1.00 | 1.00 | 1.00 | 1.00 | 1.00 | 1.00 | 1.00 | 0.75 | 0.80 | 0.33 | ---- | ---- | ---- | ---- | ---- |
| *Ldh-1* |  |  |  |  |  |  |  |  |  |  |  |  |  |  |  |  |  |  |
| 94 | 1.00 | 1.00 | 1.00 | 1.00 | 1.00 | 1.00 | 1.00 | 1.00 | 1.00 | 1.00 | 1.00 | 1.00 | 1.00 | 0.56 | 0.55 | 0.38 | ---- | ---- |
| 100 | ---- | ---- | ---- | ---- | ---- | ---- | ---- | ---- | ---- | ---- | ---- | ---- | ---- | 0.44 | 0.45 | 0.63 | 1.00 | 1.00 |
| *Mdhp-1* |  |  |  |  |  |  |  |  |  |  |  |  |  |  |  |  |  |  |
| 100 | 1.00 | 1.00 | 1.00 | 1.00 | 1.00 | 0.79 | 0.50 | 0.81 | 1.00 | 0.22 | 1.00 | 1.00 | 1.00 | 1.00 | 1.00 | 1.00 | 1.00 | 1.00 |
| 104 | ---- | ---- | ---- | ---- | ---- | 0.21 | 0.50 | 0.19 | ---- | 0.78 | ---- | ---- | ---- | ---- | ---- | ---- | ---- | ---- |
| *Icdh-2* |  |  |  |  |  |  |  |  |  |  |  |  |  |  |  |  |  |  |
| 100 | 0.94 | 1.00 | 1.00 | 1.00 | 1.00 | 1.00 | 1.00 | 1.00 | 1.00 | 1.00 | 1.00 | 1.00 | 1.00 | 1.00 | 1.00 | 0.94 | 1.00 | 1.00 |
| 110 | 0.06 | ---- | ---- | ---- | ---- | ---- | ---- | ---- | ---- | ---- | ---- | ---- | ---- | ---- | ---- | 0.06 | ---- | ---- |
| *6Pgdh* |  |  |  |  |  |  |  |  |  |  |  |  |  |  |  |  |  |  |
| 90 | 0.94 | 1.00 | 1.00 | 1.00 | 1.00 | 1.00 | 1.00 | 1.00 | 1.00 | 0.38 | 0.71 | 0.70 | 0.83 | 1.00 | 0.50 | 1.00 | 0.69 | 0.42 |
| 100 | 0.06 | ---- | ---- | ---- | ---- | ---- | ---- | ---- | ---- | 0.63 | 0.29 | 0.30 | 0.17 | ---- | 0.50 | ---- | 0.31 | 0.58 |
| *Sod-2* |  |  |  |  |  |  |  |  |  |  |  |  |  |  |  |  |  |  |
| 96 | 1.00 | 1.00 | 1.00 | 1.00 | 1.00 | 1.00 | 1.00 | 1.00 | 1.00 | 1.00 | 1.00 | 1.00 | 1.00 | 1.00 | 0.67 | ---- | ---- | ---- |
| 100 | ---- | ---- | ---- | ---- | ---- | ---- | ---- | ---- | ---- | ---- | ---- | ---- | ---- | ---- | 0.33 | 1.00 | 1.00 | 1.00 |
| *Np* |  |  |  |  |  |  |  |  |  |  |  |  |  |  |  |  |  |  |
| 100 | ---- | ---- | ---- | ---- | ---- | 0.14 | ---- | ---- | 0.18 | ---- | 1.00 | 1.00 | 1.00 | 1.00 | 1.00 | 1.00 | 1.00 | 1.00 |
| 108 | 1.00 | 1.00 | 1.00 | 1.00 | 1.00 | 0.86 | 1.00 | 1.00 | 0.82 | 1.00 | ---- | ---- | ---- | ---- | ---- | ---- | ---- | ---- |
| *Aat-1* |  |  |  |  |  |  |  |  |  |  |  |  |  |  |  |  |  |  |
| 92 | 0.71 | 0.65 | 0.79 | ---- | 0.83 | ---- | ---- | 0.31 | 0.23 | ---- | ---- | ---- | ---- | 0.29 | ---- | ---- | ---- | ---- |
| 94 | ---- | ---- | ---- | ---- | ---- | ---- | ---- | ---- | ---- | ---- | ---- | ---- | ---- | 0.50 | 0.90 | 0.38 | 0.19 | 0.18 |
| 100 | ---- | ---- | ---- | ---- | ---- | ---- | ---- | ---- | ---- | ---- | ---- | ---- | ---- | 0.07 | ---- | 0.19 | 0.38 | 0.41 |
| 102 | ---- | 0.05 | ---- | ---- | ---- | 1.00 | 1.00 | 0.63 | 0.77 | 1.00 | ---- | ---- | ---- | ---- | ---- | ---- | ---- | ---- |
| 104 | ---- | ---- | ---- | ---- | ---- | ---- | ---- | ---- | ---- | ---- | ---- | ---- | ---- | ---- | ---- | 0.44 | ---- | ---- |
| 110 | ---- | ---- | ---- | ---- | ---- | ---- | ---- | ---- | ---- | ---- | ---- | ---- | ---- | ---- | 0.10 | ---- | 0.44 | 0.41 |
| 112 | 0.29 | 0.30 | 0.21 | 1.00 | 0.17 | ---- | ---- | 0.06 | ---- | ---- | 1.00 | 1.00 | 1.00 | 0.07 | ---- | ---- | ---- | ---- |
| 117 | ---- | ---- | ---- | ---- | ---- | ---- | ---- | ---- | ---- | ---- | ---- | ---- | ---- | 0.07 | ---- | ---- | ---- | ---- |
| *Est-3* |  |  |  |  |  |  |  |  |  |  |  |  |  |  |  |  |  |  |
| 96 | 1.00 | 1.00 | 1.00 | 1.00 | 1.00 | 1.00 | 1.00 | 1.00 | 1.00 | 1.00 | 1.00 | 1.00 | 1.00 | 0.08 | 0.10 | ---- | ---- | ---- |
| 100 | ---- | ---- | ---- | ---- | ---- | ---- | ---- | ---- | ---- | ---- | ---- | ---- | ---- | 0.92 | 0.90 | 1.00 | 1.00 | 1.00 |
| *Pep-3* |  |  |  |  |  |  |  |  |  |  |  |  |  |  |  |  |  |  |
| 92 | ---- | ---- | ---- | ---- | ---- | ---- | ---- | ---- | ---- | ---- | ---- | ---- | ---- | ---- | ---- | ---- | 0.75 | ---- |
| 100 | ---- | ---- | ---- | ---- | ---- | ---- | ---- | ---- | ---- | ---- | ---- | ---- | ---- | 0.68 | 1.00 | 1.00 | 0.25 | 1.00 |
| 108 | 1.00 | 1.00 | 1.00 | 1.00 | 1.00 | 1.00 | 1.00 | 1.00 | 1.00 | 1.00 | 1.00 | 1.00 | 1.00 | 0.32 | ---- | ---- | ---- | ---- |
| *Ada-1* |  |  |  |  |  |  |  |  |  |  |  |  |  |  |  |  |  |  |
| 90 | ---- | ---- | ---- | ---- | ---- | 0.07 | ---- | ---- | 0.05 | ---- | ---- | ---- | ---- | ---- | ---- | ---- | ---- | 0.05 |
| 100 | 1.00 | 1.00 | 1.00 | 1.00 | 1.00 | 0.93 | 1.00 | 1.00 | 0.96 | 1.00 | 1.00 | 1.00 | 1.00 | 1.00 | 1.00 | 1.00 | 1.00 | 0.95 |
| *Ada-2* |  |  |  |  |  |  |  |  |  |  |  |  |  |  |  |  |  |  |
| 78 | ---- | ---- | ---- | ---- | ---- | ---- | ---- | ---- | ---- | 0.06 | ---- | ---- | ---- | ---- | ---- | ---- | ---- | ---- |
| 88 | 1.00 | 1.00 | 1.00 | 0.60 | 1.00 | 0.64 | 0.50 | 1.00 | 0.55 | 0.94 | ---- | ---- | ---- | ---- | ---- | ---- | ---- | ---- |
| 95 | ---- | ---- | ---- | 0.40 | ---- | 0.36 | 0.50 | ---- | 0.45 | ---- | 1.00 | 1.00 | 0.83 | ---- | ---- | ---- | ---- | ---- |
| 100 | ---- | ---- | ---- | ---- | ---- | ---- | ---- | ---- | ---- | ---- | ---- | ---- | 0.17 | 1.00 | 1.00 | 1.00 | 1.00 | 1.00 |
| *Gpi* |  |  |  |  |  |  |  |  |  |  |  |  |  |  |  |  |  |  |
| 100 | 0.23 | 0.35 | 0.50 | ---- | ---- | 0.07 | ---- | 0.56 | 0.55 | 0.22 | 0.29 | 0.30 | 0.67 | 0.94 | 1.00 | 1.00 | 0.88 | 0.86 |
| 108 | 0.77 | 0.65 | 0.50 | 1.00 | 1.00 | 0.93 | 1.00 | 0.44 | 0.46 | 0.78 | 0.71 | 0.70 | 0.33 | 0.06 | ---- | ---- | 0.13 | 0.14 |
| *Pgm-1* |  |  |  |  |  |  |  |  |  |  |  |  |  |  |  |  |  |  |
| 100 | 1.00 | 1.00 | 1.00 | 1.00 | 1.00 | 1.00 | 1.00 | 1.00 | 1.00 | 1.00 | 1.00 | 1.00 | 1.00 | 1.00 | 0.30 | 1.00 | 1.00 | 1.00 |
| 106 | ---- | ---- | ---- | ---- | ---- | ---- | ---- | ---- | ---- | ---- | ---- | ---- | ---- | ---- | 0.70 | ---- | ---- | ---- |
| *Pgm-2* |  |  |  |  |  |  |  |  |  |  |  |  |  |  |  |  |  |  |
| 100 | 1.00 | 0.90 | 1.00 | 1.00 | 1.00 | 1.00 | 1.00 | 1.00 | 1.00 | 1.00 | 1.00 | 1.00 | 1.00 | 1.00 | 0.30 | 1.00 | 1.00 | 1.00 |
| 110 | ---- | 0.10 | ---- | ---- | ---- | ---- | ---- | ---- | ---- | ---- | ---- | ---- | ---- | ---- | 0.70 | ---- | ---- | ---- |
| **Variability** |  |  |  |  |  |  |  |  |  |  |  |  |  |  |  |  |  |  |
| $H_E$ | 0.05 (0.03) | | | | | 0.09 (0.03) | | | | | 0.06 (0.03) | | | 0.14 (0.04) | | 0.08 (0.04) | | |
| $H_O$ | 0.04 (0.02) | | | | | 0.06 (0.02) | | | | | 0.07 (0.04) | | | 0.10 (0.04) | | 0.05 (0.03) | | |
| $A_R$ | 1.2 (0.4) | | | | | 1.3 (0.5) | | | | | 1.2 (0.5) | | | 1.6 (1.0) | | 1.3 (0.5) | | |

The spatially explicit Bayesian clustering analyses conducted with TESS based on allozyme variation, indicated K=5 as the best grouping option for this dataset. Indeed, higher values of K yielded only minimal variation in values of the DIC statistic (Figure 2B), and did not turn into biologically interpretable results (not shown).With K=2 (Figure 2C), individuals were grouped into a peninsular cluster (samples 1-14), and an alpine cluster (samples 18-21), thus fully matching the geographic distribution of the Italian endemic subspecies *S. s. gigliolii* and the nominal subspecies *S. s. salamandra,* respectively. The geographically intermediate sample 16 was the only one showing substantial evidence of admixture between both groups. When the best clustering option for K=3 was considered (Figure 2D), peninsular samples were further subdivided into two groups with negligible evidence for admixture (see sample 9 in Figure 2D). One group clustered southern samples (1-10), while the other clustered northern peninsular samples (12-14). With the best clustering option for K=4 (Figure 2E), southern samples were further allocated to two distinct groups arranged along the north-south axis. In this case, evidence for admixture were substantial and appeared mostly asymmetric, from south to north. Finally, with K=5 (Figure 2F) samples drawn from the alpine arc (19-21) were assigned to a distinct cluster. Among them, sample19 (i.e. the one in closer geographic contiguity to the remaining samples), was the only one showing evidence of mixed ancestry.

**Figure 2.**
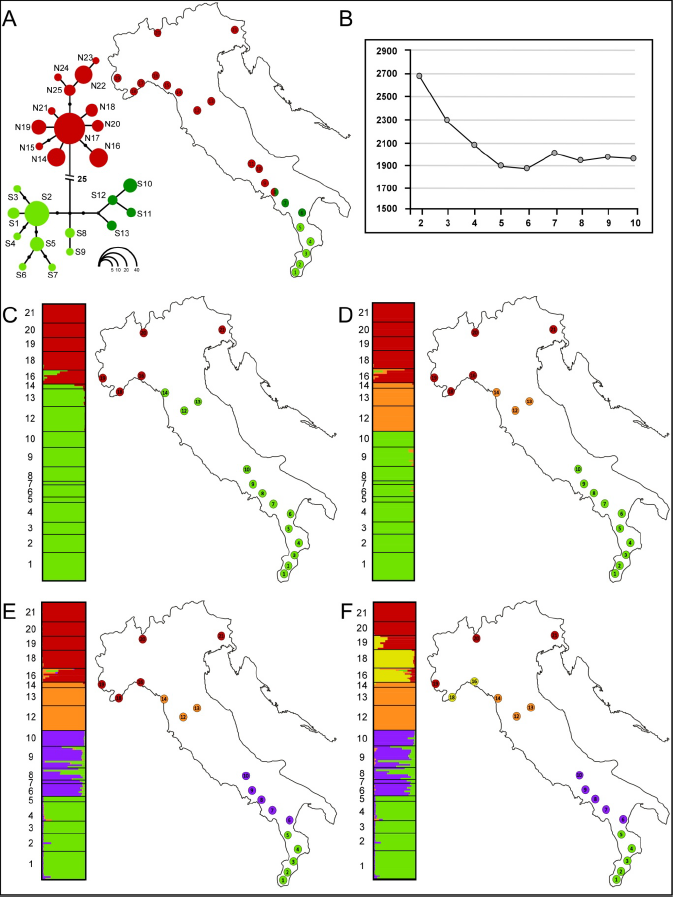
Population genetic structure of *S. salamandra* in Italy. (A) mtDNA haplotype genealogy generated using HaplotypeViewer, based on the ML phylogenetic tree, and geographic distribution of the main haplotype groups. Circle sizes are proportional to haplotype frequency (see inset, lower right), whereas missing intermediate haplotypes are shown as dots. Population samples are coloured as pie diagrams, showing the geographic distribution of the main haplogroups. (B) Mean values of the DIC statistic (averaged over 10 runs) estimated for models with K ranging from 2 to 9. (C)-(F) Results of the Bayesian clustering analyses carried out with TESS and BAPS for values of K between 2 and 5. Bar-plots show individual admixture proportions for the genetic clusters inferred using TESS. Populations assigned to the same cluster by BAPS are marked by distinct colours on the maps. Maps and diagrams were drawn using the software Canvas 11 (ACD Systems of America, Inc.).

The sample-based Bayesian clustering method implemented in BAPS, yielded results fully consistent with those obtained with TESS, for all values of K between 2 and 5 (Figure 2C-2F). Nevertheless, it suggested K=7 as the best clustering option. On the other hand, both with K=6 and K=7 single populations were assigned to distinct clusters. Identical results were obtained running this analysis either with or without using geographic location of samples as prior information.

Estimates of genetic diversity were computed for sample pools consistent with K=5, that is, the highest level of population structure identified by both TESS and BAPS analyses. As shown in Table 2,the lowest level of diversity was observed at all parameters for the southernmost group (pooling samples 1-5), whereas the highest values were observed for the north-western group (pooling samples 16-18).

The final mtDNA sequence alignment comprised 1220 bp for all the 158 *S. salamandra* individuals analysed. The cytochrome B gene fragment analysed (hereon CYTB) was 582 bp in length, with 23 variable positions of which 21 were parsimony informative and the cytochrome oxidase subunit I gene fragment (hereon COI) was 638 bpin length, and showed29 variable positions, of which 24 were parsimony informative. Twenty-five unique haplotypes were identified in the concatenated dataset, whose distribution among the sampled populations is shown in Table 1.

The mtDNA haplotype genealogy estimated using HaplotypeViewer is presented in Figure 2A. Two main groups of closely related haplotypes were observed, separated from one another by 25 mutational steps. One haplogroup (green in Figure 2A; haplotype series S) was geographically restricted to the south of the Italian peninsula (samples 1-8), whereas the other haplogroup (red in Figure 2A; haplotype series N) was widespread from the alpine arc to samples in south-central Italy (samples 8-21). Syntopy between both haplogroups was only found at the geographically intermediate sample 8. Within the southern haplogroup, to sub-groups of haplotypes were found, one restricted to samples from the Calabria region, and the other restricted to more northern samples (dark and light green in Fig. 2A, respectively). Syntopy between both sub-groups was not observed.

## Discussion

Our analyses of mitochondrial and nuclear genetic variation concordantly, and not unexpectedly (see Introduction), show that two main lineages of *S. salamandra* occur along the Italian peninsula. However, this is the only line of concordance observed among the patterns of variation of both marker classes. First, populations marking the geographic transition between groups have been observed both with mitochondrial and with nuclear markers (samples 8 and 16), but the respective geographic locations were more than 600 km apart (see Fig. 2A and 2C). Secondly, mitonuclear discordances also emerged at shallower levels of population structure. The highest number of discernible geographic sub-groups, with nuclear and mitochondrial markers respectively, was five and three (Fig. 2A and 2F). Thirdly, an imprint of asymmetric gene flow (from south to north) was observed between two sub-groups of the southern lineage at the nuclear dataset (green and purple sub-groups in Fig. 2E and 2F), whereas no imprints of admixture was observed between them at the mitochondrial dataset (light green and dark green in Fig. 2A).

Mitonuclear discordances are not uncommon in amphibians (e.g.^4, 27–30^), but the extent of discordance observed in the present study is conspicuous. The geographic displacement between mitochondrial and nuclear contact zones among the two main lineages, largely exceeds the extent of discordances previously noted among fire salamander lineages from different geographic regions^21–23^, and has no parallels among different taxa from peninsular Italy. Interestingly, both mtDNA and nuclear contact zones are located within well known suture zones (sensu^31^), where interspecific and intraspecific hybrid zones and range edges have been previously reported for various taxa, including amphibians (e.g.^4, 32–35^). Therefore, lines of cross-taxon concordance cannot be used here as inferential clues in the attempt to reconcile such discordance into a population history. Even a comparison of genetic and morphological patterns of variation appears uselessin this regard. In fact, the closer concordance of morphological variation with allozyme variation, rather than with mtDNA, might be accounted for either by a primary location of the contact zone in the north-western Apennines, or by a northward migration of the contact zone, with displacement of northern traits by the advancing southern lineage.

The close affinity among mtDNA sequences of individuals sampled in northern Balkans, northern Europe, southern Alps, and north-western Apennines (this study and^24^), suggests that the northern lineage (namely *S. s. salamandra)* colonized the Apennines as the last step of a large-scale range expansion out of the Balkans into Western Europe^24^. Here, it underwent a secondary contact with the southern, peninsular endemic lineage (namely *S. s. gigliolii),* most likely during the post-glacial epoch^24^. But where did the two lineages first meet and mate? And what processes might best explain the observed mito-nuclear discordance? At least two conflicting scenarios could be delineated in this respect. The first that the secondary contact between both lineages first occurred in the north-western Apennines, as depicted by nuclear data (Fig. 2C), and that this event was followed by rampant introgression of the *S. s. salamandra* mitochondrial lineage into the *S. s. gigliolii* populations in northern and central Apennines. The second that the secondary contact occurred in south-central Apennines, as depicted by the mtDNA transition, and that the hybrid zone between nuclear genomes has moved northward to its current location, not followed by a movement of the mtDNA contact zone. Both rampant mitochondrial introgression and hybrid zone movement have been repeatedly invoked to explain instances of mitonuclear discordances^7^, and several underlying mechanisms have been proposed^7, 8, 10, 36, 37^, but in the absence of replicated temporal samples or of replicated contact zones at disparate sites, disentangling these competing scenarios is a challenging task, since most of the genetic patterns of variation expected under one scenario do not entirely exclude the other^38^. In the present case, however, two main lines of evidence lead us to favour rampant mtDNA introgression over a hybrid zone movement from south to north.

A first useful indication in this regard comes from the spatial genetic structure observed with allozymes. Indeed, in a moving hybrid zone scenario, a unidirectional clinal tail of admixture is expected, as a consequence of the movement (^38^and references therein). Our scattered sampling and, more generally, the highly fragmented geographic distribution of the fire salamander through north-central and central Apennines, prevented us from carrying out a formal and straightforward cline analysis. Nonetheless, the strongly structured pattern of genetic variation that emerged with the Bayesian clustering analyses, does not conform with this expectation. In fact, it showed the occurrence of genetic clusters along the north-south axis, and a lack of clines in admixture coefficients within the putative area of hybrid zone movement, at all the hierarchical levels of the analysis. Furthermore, not even the geographic variation observed at the level of the single allozyme loci shows clear and concordant patterns of intergradation.

A second indication comes from the Upper Pleistocene fossil record. A scenario of post-glacial colonization of northern and central Apennines by *S. s. salamandra,* followed by the establishment of a hybrid zone in south-central Apennine and its subsequent displacement northward to its current location, would imply the absence of fire salamanders in the northern Apennines until the post-glacial arrival of *S. s. salamandra.* However, this scenario is in sharp contrast with a fire salamander fossil record found in north-western Apennines (Grotta di Equi^39^), close to our sampling site 14, and dated back to the last glacial epoch (around 45 000 years bp). The occurrence of this fossil record is instead fully consistent with the alternative scenario of a secondary contact zone primarily located in the north-western Apennines, and a rampant mtDNA introgression southward. Under this scenario, the genetic cluster emerging at K=3, grouping samples 12-14, would reflect the presence of *S. s. gigliolii* in this area during the last glaciation, and the occurrence of a glacial sub-refugium within the region, as already suggested for several temperate species (see^35^ and references therein).

Under the scenario of rampant mtDNA introgression, then the main question to answer is, why did *S. s. salamandra* mtDNA introgressed so deeply into *S. s. gigliolii* populations? Providing a definitive answer goes behind the scope of this study, and would require specific experimental designs. However, it is worth noting that several of the most commonly invoked causes of mito-nuclear discordance can already be dismissed in the present case, allowing us to substantially narrow future research efforts around a restricted set of putative causes. First, sex-biased processes, such as female-biased dispersal^40^ or disassortative preferences in female mate choice^30^, are unlikely to have played a role. In fact, given the modest dispersal abilities of *S. salamandra^41^,* these processes would have had take active for many generations in order to allow the uniparentally inherited mtDNA wake to travel more than 600 km to the south, without leaving imprints at the nuclear genome. Moreover, although extensive surveys are still lacking, the available evidence does not support female biased dispersal in *S. salamandra^41^.* Second, purely stochastic processes related to demographic disparities between interactive lineages upon secondary contact^8, 36^, are unlikely to have been implicated either. Literature surveys and simulation-based works^7, 8^ ^36^ strongly suggest that differential introgression triggered by these processes is more likely to occur at the uniparentally inherited markers, and from the resident toward the expanding lineage. In the present case, this pattern would imply massive introgression of mtDNA from east to west of the contact zone, that is, in the opposite direction actually observed. In this regard, it is also worth mentioning that our analyses cast doubts on the hypothesized long-distance range expansion of *S. s. salamandra,* from the Balkan into the Apennine chain. Indeed, at the shallower level of the hierarchical population structure, clustering analyses suggest the occurrence of a differentiated cluster in north-western Italy (Fig. 2F), whereas haplotype sharing between populations sampled in north-west (samples 17-19) and the Alps (samples 20-21) was not observed (see Table 1). In turn, this indication of a shallow genetic structure within the *S. s. salamandra* lineage in western Italy suggests that its eastward range expansion might have begun close to the most north-eastern cluster of *S. s. gigliolii,* as already observed in the alpine newt and several other taxa within the same area (see^35^ and references therein). In turn, this scenario would further weaken the plausibility of a major demographic disparity between lineages at the time of formation of the secondary contact zone, since (loosely speaking) the secondary contact would have occurred between two “residents”, rather than between one resident and one expanding lineage. Consequently, an adaptive introgression of the *S. s. salamandra* mtDNA into *S. s. gigliolii* appears the most plausible explanation for the observed mitonuclear discordance. Of course, given the many organismal functions in which the mitochondrial genome is implicated^42–45^, and the unsuitability of our data in this respect, we refrain from even hypothesizing which adaptive advantage promoted the massive mtDNA introgression. In light of the frequently observed mitonuclear discordances in *S. salamandra,* at various levels of population structure^21–23^, we see this issue as a major research question, amenable to future experimental efforts.

Finally, in addition to the major mitonuclear discordance discussed above, another instance of discordant patterns of gene exchange, emerged also at the shallower level of population structure represented by the two sub-gropus identified within the southern lineage (Fig. 2). In this case, the structure of the discordance does not match the major one discussed above, as mtDNA and allozyme transition zones overlap (see Fig. 2A and 2E). However, while no evidence of admixture was apparent at the mtDNA, substantial and asymmetrical introgression (from south to north) was observed with allozymes. This geographic pattern of variation closely mirror those observed within several contact zones between differentiated lineages of fire salamander in the Iberian peninsula^21, 23^. Nuclear mediated gene-flow, driven by male-biased dispersal across the contact zone, or by the competitive advantage of males of one lineage over the other, might comfortably explain these geographic patterns (see also^30^). However, as already mentioned above, sex-biased dispersal has not been observed in *S. salamandra* populations^41^ and, as pointed out by Pereira, Martinez Solano & Buckley^23^, we still lack fundamental knowledge of the mechanisms underlying the observed (and recurrent) patterns. Besides going deeper into the genomic patterns of geographic variation, future studies aimed at shedding light on these mechanisms, will have to gather and conjugate information across the secondary contact zones about variation of life-history traits and the multivariate phenotype, hopefully taking advantage of a comparative trans-geographic experimental approach.

## Methods

### Sampling

We collected 188 individuals of *S. salamandra* from 21 localities, ranging from the southern slope of the Alps to the southern tip of the Italian peninsula (Figure 1 and Table 1). Each individual was anaesthetized in the field by submersion in a 0.02% solution of MS222 (3-aminobenzoic acid ethyl ester), and tissue samples were taken through a toe-clipping procedure, then each individual was released in its sampling point. Collected tissues were brought to laboratory in liquid nitrogen, and then stored at − 80 °C. Field works and collection of tissues were approved by the Italian Ministry of Environment (permit numbers: DPN-2009-0026530) and were performed in accordance with the relevant guidelines and regulations.

### Laboratory procedures

Standard horizontal starch gel (10%) electrophoresis was used to investigate the genetic variation of 23 putative allozyme loci. Enzyme systems analyzed, putative loci, and buffer systems used in electrophoretic procedures are listed in Table 3. Alleles were called by their mobility (in mm) with respect to the to the most common (100) in a reference population (Campone).

**Table 3.** Enzymes systems analysed in *S. salamandra* (EC: Enzyme Commission number), encoding loci, and buffer systems used for the allozyme electrophoresis procedure.

| Enzyme | EC | Encoding loci | Buffer systems |
| --- | --- | --- | --- |
| Glycerol-3-phosphate dehydrogenase | 1.1.1.8 | G3pdh | 5 |
| Lactatedehydrogenase | 1.1.1.27 | Ldh-1, Ldh-2 | 4 |
| Malate dehydrogenase | 1.1.1.37 | Mdh-1, Mdh-2 | 5 |
| Malate dehydrogenase (NADP+) | 1.1.1.40 | Mdhp-1, Mdhp-2 | 2,5 |
| Isocitratedehydrogenase | 1.1.1.42 | Icdh-1, Icdh-2 | 6 |
| 6-Phosphogluconate dehydrogenase | 1.1.1.43 | 6Pgdh | 5 |
| Superoxidedismutase | 1.15.1.1 | Sod-1, Sod-2 | 3 |
| Purine-nucleoside phosphorylase | 2.4.2.1 | Np | 3 |
| Aspartatetransaminase | 2.6.1.1 | Aat-1, Aat-2 | 5 |
| Carboxylesterase | 3.1.1.1 | Est-3 | 1 |
| L-LeucylLeucylLeucinePeptidase | 3.4.11 | Pep-3 | 2 |
| Adenosine deaminase | 3.5.4.4 | Ada-1, Ada-2 | 2 |
| Mannosephosphateisomerase | 5.3.1.8 | Mpi | 3 |
| Glucosephosphateisomerase | 5.3.1.9 | Gpi | 3 |
| Phosphoglucomutase | 5.4.2.2 | Pgm-1, Pgm-2 | 4 |

Buffer systems: 1) DiscontinuousTris/CitratepH 8.7<sup>63</sup>; 2) ContinousTris/CitratepH 8.0<sup>64</sup>; 3) Tris/Versene/BoratepH 8.0<sup>65</sup>; 4) Tris/MaleatepH 7.4<sup>65</sup>; 5) Phosfate-CytratepH 6.3<sup>66</sup>.

Genomic DNA was extracted using proteinase K digestion followed by a standard phenol-chloroform protocol^46^. Polymerase chain reactions (PCR) were carried out to amplify portions of the two mitochondrial gene fragmentsCYTBand COI.After preliminary PCR amplifications using generic primers drawn from the literature^47, 48^, more specific primer pairs were designed and used to amplify and sequence all the individuals analysed. Primers designed and used to amplify the CYTB gene fragment were 494-salamod-CATCAACATCTCCTACTGATGAAA and CYTB-salamod-GGAGTAAGC AGTGAGATTAC C, whereas VF1d-TTCTCAACCAACCACAARGAYATYGGand VR1d-TAGACTTCTGGGTGGCCRAARAAYCA^49^were used to screen variation at the COI gene fragment. Amplifications were carried out in a final volume of15 μl containing: MgCl2 (2 mM), four dNTPs (0.2mM each), primers (0.2 μM each), the enzyme Taq polymerase (0.5 U, Promega), reaction buffer (1X, Promega) and 2 μl of 20 ng/μl DNA template. PCR cycling conditions were the same for both the genes: a step at 94°C for 5 min followed by 36 cycles of 45 sec at 94°C, 1 min at 55°C (CYTB) or 51°C (COI), and 1.5 min at 72°C, followed by a single final step at 72°C lasting 10 min. The PCR products were purified and sequenced by Macrogen Inc. (http://www.macrogen.com). All sequences were deposited in GenBank (accession numbers: XXXX-XXXX).

### AUozymes data analysis

Allele frequencies and genetic diversity estimates, including allelic richness, observed heterozygosity, and unbiased expected heterozygosity^50^, were estimated using the programs GENETIX 4.05.2 software^51^. The occurrence of Hardy-Weinberg and genotypic linkage equilibriawere addressed for each locus and locus-pair, respectively, in each sample by means of exact tests as implemented in FSTAT 2.9.3^52^.

We investigated the genetic structure of *S. salamandra* populations along peninsular Italy, using two Bayesian clustering methods: the individual-based and spatially explicit approach implemented by TESS 2.3.1^53^, and sample-based approach implemented in BAPS v 6^54^.

The analysis with TESS was carried using the admixture model, the option to update the spatial interaction parameter activated, and all other settings left to the default options. A preliminary analysis was conducted to restrict the range of plausible K values (i.e. the number of clusters). Values of K between 1 and 21 were tested, with 10 replicates per K, each of 5000 iterations following 3000 iterations discarded as burn-in. The settings for the final run were: K between 2 and 10, 100 replicates per K, and a burn-in of 30 000 iterations followed by 50 000 iterations. For each value of K, the 10 replicates with the lowest Deviance Information Criterion (DIC) were permuted in the software CLUMPP 1.1.2^55^, and the resulting clustering was visualized with DISTRUCT^56^.

The Bayesian clustering analysis with BAPS was carried out using population samples as the units of the analysis. With this method we run two sets of analyses, with and without using geographic coordinates as prior information to infer the best number of clusters (K). In both cases, we explored values of maximal K between 2 and 10, carrying out 5 replicates for each maximal K to check for consistency among runs.

When interpreting results from both TESS and BAPS analyses, we followed recommendations by Meirmans^57^, that is, we presented and discussed results of each clustering option deserving biological interpretation, irrespective of the K-value that is deemed ‘optimal’ according to the summary statistics.

### Mitochondrial DNA data analysis

Sequence electropherograms were controlled by eye using the software FinchTv1.4.0 (Geospiza Inc.), and the sequence alignments were obtained using CLUSTALX 2.0^58^.Sequences of the two gene fragments analysed were concatenated using the software CONCATENATOR 1.1.0^59^. The best partitioning strategy for the concatenated mtDNA dataset was assessed by means of the software PartitionFinder v1.0.1^60^, using the Bayesian information (BI) criterion and the ‘greedy’ search method. This analysis suggested that the HKY substitution model best fitted all the data partition considered (1^st^, 2^nd^, and 3^rd^ position for either COI or CYTB fragments).

The phylogenetic relationships between haplotypes were inferred by means of the maximum likelihood algorithm as implemented in PhyML 3.10^61^, using default settings for all parameters but the substitution model (HKY) and the type of tree improvement (SPR and NNI). The estimated tree was then converted into an haplotype network using the software HaplotypeViewer^62^.

The overall pattern of genetic variation observed at level of the entire dataset, suggested that the mtDNA variation might be not entirely reflective of the population structure and history of *S. salamandra* in Italy (see Results). Consequently, we refrained from using mtDNA variation to estimating historical demographic processes, phylogeographic dynamics, and their time line.

## Acknowledgements

We dedicate this paper to the loving memory of our friend and colleague Francesco Spallone. We are grateful to Gaetano Aloise, Claudio Bagnoli and Florinda Sacco for their help with sample collection and/or with laboratory procedures. This work was supported by a grant from the Italian Ministry of Education, University, and Research (PRIN project 2012FRHYRA).

## Author contributions statement

DC and GN conceived and designed the study. RB and PA performed the experiments. RB, DP and DC analysed the data. RB and DC wrote the paper. All authors reviewed the manuscript.

## Additional information

### Accession codes

Genbank accession numbers: XXXX-XXXX (to be populated upon acceptance).

